# BLOSUM Is All You Learn Generative Antibody Models Reflect Evolutionary Priors

**DOI:** 10.1101/2025.10.26.684652

**Authors:** Talip Uçar, Pietro Sormanni

## Abstract

Generative models have emerged as powerful tools for antibody sequence design, with recent studies demonstrating that log-likelihood scores from these models can correlate with binding affinity and potentially serve as effective ranking metrics. This raises a fundamental question: why should log-likelihood scores from generative models correlate with binding affinity? In this work, we investigate the biochemical basis of these model-derived log-likelihoods by comparing them with classical evolutionary similarity metrics. We find that BLOSUM similarity scores between designed and parental antibody sequences correlate strongly with measured binding affinity—on par with the predictive performance of a state-of-the-art diffusion-based generative model. Moreover, these BLOSUM scores also align closely with log-likelihoods from multiple generative models, suggesting that such models may be implicitly learning evolutionary priors encoded in substitution matrices. When computed with respect to a known binder, both BLO-SUM scores and log-likelihoods act as approximate measures of sequence distance from that reference. As this distance increases, the likelihood of a candidate being a binder decreases, explaining the observed correlation between these scores and binding affinity. In contrast, similarity scores based on position weight matrices (PWMs) and position-specific scoring matrices (PSSMs), which do not rely on knowledge of the parental sequence, show weaker and less consistent alignment with binding affinity, with performance depending on the background sequence data. Additionally, using consensus sequences in place of parental sequences to compute BLOSUM scores largely eliminates the observed correlation with affinity, underscoring the context-specific nature of the correlations. These findings highlight the potential of interpretable, evolution-inspired metrics to complement generative modeling in antibody design, offering insights into both model behavior and biological relevance.

## 1. Introduction

Antibody design continues to be a critical area of research for both therapeutic and diagnostic applications, where the goal is to target antibodies toward a predetermined epitope of interest, optimize binding affinity and specificity, and improve overall developability of antibody candidates. Computational methods, particularly generative models trained on large-scale antibody sequence datasets, have shown significant promise in accelerating this process. These models can propose candidate sequences that adhere to learned patterns of natural antibody repertoires, potentially capturing features relevant to biological function and molecular recognition. A major challenge, however, lies in the development of reliable in silico metrics to rank designed sequences by their likelihood of exhibiting high binding affinity.

Among available metrics, log-likelihood scores derived from generative models trained solely on structural and/or sequence data have recently been shown to correlate with experimentally measured binding affinity (Ucar et al., 2024), indicating their potential as a practical tool for prioritization. Yet the biochemical basis of this correlation remains poorly understood. It is not clear whether these scores reflect meaningful biophysical or evolutionary constraints, or whether they serve merely as model-specific heuristics.

In this work, we investigate the biochemical grounding of model-derived scores by examining their relationship with classical sequence similarity metrics. Specifically, we assess whether these scores align with metrics grounded in evolutionary substitution dynamics, including BLOcks SUbstitution Matrix (BLOSUM) similarity (Henikoff & Henikoff, 1992), position weight matrices (PWM) (Stormo et al., 1982), and position-specific scoring matrices (PSSM) (Gribskov et al., 1987). These metrics are computed using different strategies: BLOSUM similarity is evaluated between designed sequences and various types of reference sequences, while PWM and PSSM scores are derived from amino acid frequency profiles obtained from large antibody datasets.

By comparing these scores with both measured binding affinities and log-likelihoods from multiple generative models—including a diffusion-based model DiffAbXL-A (Ucar et al., 2024), an inverse folding model AntiFold (Høie et al., 2023), and a language model IgLM (Shuai et al., 2023), we aim to clarify whether such models capture biologically meaningful substitution patterns. Our results highlight the importance of context-specific similarity metrics and suggest that interpretable, evolution-inspired scores can complement generative models by improving the transparency and reliability of model-guided antibody design pipelines.

## 2. Related Work

Generative models have become increasingly important in protein and antibody design, leveraging advancements in deep learning to generate novel sequences with desirable properties. These models typically fall into three broad categories: large language models (LLMs) trained on sequence data (Malherbe & Uçar, 2024), graph-based models that capture spatial and topological properties (Kong et al., 2023), and diffusion-based models that simulate denoising processes over sequence and/or structure spaces (Ucar et al., 2024). Each approach differs in how it represents and generates protein information—ranging from purely sequence-based generation to more complex structure-aware co-design frameworks.

While much of the recent focus has been on improving generative quality and integrating structure prediction tools (e.g., AlphaFold (Jumper et al., 2021), RoseTTAFold (Baek et al., 2021)), a parallel challenge lies in evaluating and ranking generated designs. Commonly used in silico metrics include structural scores (e.g., RMSD, ipTM, pAE) (Abramson et al., 2024) and sequence-based scores such as amino acid recovery (AAR) or log-likelihood under pretrained models (Ucar et al., 2024; Luo et al., 2022). However, these metrics are not explicitly optimized to reflect functional outcomes like binding affinity, which limits their utility in candidate prioritization.

Recent studies have begun to explore whether log-likelihood scores from generative models can serve as more functionally meaningful evaluation metrics. For instance, Shanehsaz-zadeh et al. (2023b) observed that higher-likelihood sequences generated by IgMPNN yielded higher proportions of binders, though the correlation was indirect and assessed via enrichment metrics. Other work in general protein fitness prediction has used zero-shot likelihood ranking across mutational scans (Truong Jr & Bepler, 2023), but the results have varied across assay types. Specifically for antibodies, Chungyoun et al. (2024) found that log-likelihood does not always align with functional readouts such as binding or expression, suggesting that model scores alone may be insufficient.

More recently, Ucar et al. (2024) systematically evaluated log-likelihood scores from a diffusion-based antibody generative model (DiffAbXL-A), finding that these scores consistently and significantly correlate with measured binding affinity across diverse datasets. This supports the view that generative models can internalize biophysical and evolutionary constraints relevant to antigen binding, even without explicit supervision on affinity labels. Moreover, BLO-SUM matrices have long been used to quantify functional similarity between sequences and are known to reflect evolutionary constraints. Prior efforts have used such metrics for sequence alignment and clustering but not explicitly for affinity ranking in design contexts (Altschul et al., 1990). BLOSUM scores have also been used to estimate mutation effects on binding stability via NQFEP simulations (Gessner et al., 2024), without direct comparison to experimental affinity. This work builds on prior studies by directly investigating the biochemical basis of log-likelihood as a scoring function. We examine whether classical evolutionary similarity metrics—such as those derived from BLOSUM, PWM, and PSSM—are predictive of binding affinity and whether they correlate with the scores produced by generative models.

## 3. Method

We evaluate the relationship between log-likelihood scores from generative models and classical evolutionary metrics for predicting antibody binding affinity. Specifically, we compare four approaches: (i) log-likelihoods from three representative generative models — a diffusion-based model, an inverse folding model, and a language model (Ucar et al., 2024; Høie et al., 2023; Shuai et al., 2023), (ii) BLOSUM-based scoring (Henikoff & Henikoff, 1992), (iii) PWM–derived scores (Stormo et al., 1982), and (iv) PSSM–derived scores (Gribskov et al., 1987). All methods are evaluated across multiple datasets with experimentally measured affinities.

### Scoring Region Masking

To ensure consistent comparison across scoring methods, we define a dataset-specific mutation mask ℳ that identifies positions relevant for scoring. For BLOSUM, PWM, and PSSM-based evaluations, we first align all sequences using the AHo numbering scheme (Honegger & PluÈckthun, 2001), and include a position in ℳ if it differs between the parental sequence (i.e., the wild type, WT) and any of its variants. In contrast, for log-likelihood–based scoring using the DiffAbXL-A and IgLM models, no alignment is performed; the mask is computed directly from raw sequence positions that differ from the parental sequence. For AntiFold, which uses IMGT numbering internally to define complementarity-determining regions (CDRs), we provide the model with the specific list of CDRs designed in each library to define ℳ. This approach maintains compatibility with how these generative models process input and ensures that each method is evaluated in its appropriate context.

### Log-likelihood scoring

We evaluate log-likelihoods using three different generative models: DiffAbXL-A, AntiFold, and IgLM.

#### DiffAbXL-A

DiffAbXL-A is a scaled variant of the diffusion-based model DiffAb (Luo et al., 2022), trained to generate all six complementarity-determining regions (CDRs) of an antibody given structural context. The model is trained on an expanded synthetic dataset with longer input lengths, improving its generalization ability (Ucar et al., 2024). Log-likelihoods are computed in **De Novo (DN)** mode, where both sequence and structure are masked over the scoring region. We refer to this mode as DiffAbXL-A-DN. For a designed sequence *s*, and positions *j* in the mutation mask ℳ, we define the score:

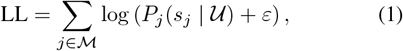

where *P*_*j*_(*s*_*j*_| 𝒰) is the probability of residue *s*_*j*_ predicted by the model given unmasked context 𝒰, and *ε* is a small constant (e.g., 10^−9^) for numerical stability.

#### AntiFold

AntiFold is an inverse folding model based on ESM-IF1 (Høie et al., 2023), which autoregressively predicts sequence from structure. To compute log-likelihoods, we first pass either the parental or mutant sequence along with its backbone structure into the model to obtain per-position log-probabilities. We then gather the log-probabilities corresponding to the mutant amino acids at the mutated positions ℳ. In the main analysis, we use the parental sequence for context (**AntiFold**_PA_). The log-likelihood is defined as:

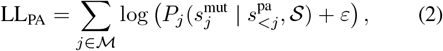

where 𝒮 is the structure, 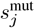 is the mutant residue at position *j*, and 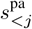 is the prefix of the parental sequence.

#### IgLM

IgLM is a decoder-only language model trained with a masked span infilling objective on antibody sequences (Shuai et al., 2023). We evaluate log-likelihood using two scoring modes: preceding context only ([pre]) and bidirectional context ([bi]). In both cases, we use the parental sequence as context in the main experiments. Mutation regions may consist of one or more contiguous spans (e.g., individual or multiple CDRs), each of which is scored independently in bidirectional context case.

##### Preceding context only ([pre])

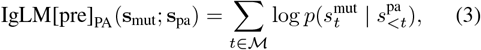

where the parental sequence **s**_pa_ is passed to the model to obtain logits, and the mutant sequence **s**_mut_ is used to select log-probabilities at mutation sites *t* ∈ ℳ.

##### Bidirectional context ([bi])

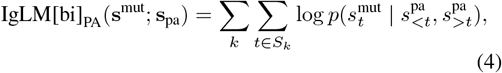

where each mutated span *S*_*k*_ is a contiguous region (e.g., a CDR) that is masked in the parental sequence, and the model predicts the mutant residues using bidirectional context. Multiple spans can be masked and evaluated independently if the mutation mask covers disjoint regions. Furthermore, since IgLM is trained to model masked spans using bidirectional context, scoring based on bidirectional information is better aligned with its training objective and has been empirically shown to produce lower perplexity than scoring based solely on preceding context (Shuai et al., 2023).

Results for scoring using the mutant sequence as context (IgLM[pre]_MUT_, IgLM[bi]_MUT_, and AntiFold_MUT_), along with additional details on how the scores are computed, are provided in Sections C - F of the Appendix.

### BLOSUM similarity scoring

BLOSUM similarity is computed using substitution matrices (e.g., BLOSUM45, 62, 80, and 90). For each designed sequence, similarity is calculated with respect to one of the following reference sequences: (1) the parental sequence from the same dataset (denoted as BLOSUM_PA_); (2) a global consensus sequence derived from two antibody datasets—either human antibodies from the Observed Antibody Space (OAS) or antibody-antigen complexes from SAbDab—denoted as 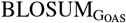 and 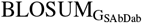, respectively; or (3) a consensus sequence derived from the antibody variants within each dataset, referred to as dataset-specific (DS) and denoted as BLOSUM_DS_. For (2), separate consensus sequences are constructed for heavy, kappa, and lambda chains using AHo numbering (Honegger & PluÈckthun, 2001). The similarity score is defined as:

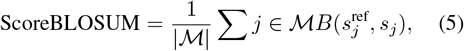

where 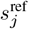 is the residue at position *j* in the reference sequence, *s*_*j*_ is the corresponding residue in the designed sequence, and *B*(*a, b*) is the substitution score from the chosen BLOSUM matrix.

### PWM-based similarity scoring

Position weight matrices (PWMs) are constructed from aligned human antibody sequences in the OAS and SAbDab databases. Sequences are split by chain type (heavy, kappa, and lambda) and aligned using AHo numbering. From these alignments, we compute amino acid frequency distributions at each position, producing normalized matrices in which each column sums to 1.

For each designed sequence, we compute the PWM score as the sum of amino acid frequencies at the mutated positions, matched by chain type:

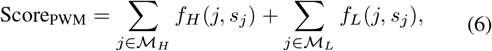

where *f*_*H*_ (*j, s*_*j*_) and *f*_*L*_(*j, s*_*j*_) are the amino acid frequencies at position *j* in the heavy and light chain PWMs, respectively, and ℳ_*H*_, ℳ_*L*_ are the subsets of ℳ corresponding to the heavy and light chains.

### PSSM-based similarity scoring

Position-specific scoring matrices (PSSMs) are computed from aligned antibody sequences, where each entry reflects the log-odds score of observing amino acid *a* at position *j* relative to background. We derive PSSMs from three sources: (1) human antibodies from the OAS repertoire, (2) the SAbDab database, and (3) the sequences within each experimental dataset, where a separate PSSM is constructed for each dataset using only its constituent sequences. We refer to this third approach as dataset-specific PSSMs (**PSSM**_**DS**_). As with PWMs, sequences are aligned using AHo numbering and split by chain type.

For each designed sequence, we compute the PSSM score as:

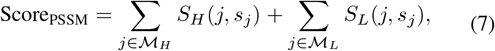

where *S*_*H*_ (*j, s*_*j*_) and *S*_*L*_(*j, s*_*j*_) denote the log-odds substitution scores from the PSSMs for the heavy and light chains, respectively. Higher scores indicate higher evolutionary preference for the observed amino acids at those positions. Additional details on the construction of PSSMs and PWMs can be found in Section B of the Appendix.

### Hydrophobicity and rigidity scoring

To evaluate coarsegrained biophysical trends, we compute the average hydrophobicity and rigidity of mutated residues using the Kyte-Doolittle (Kyte & Doolittle, 1982) and Karplus-Schulz (Karplus & Schulz, 1985) scales, respectively:

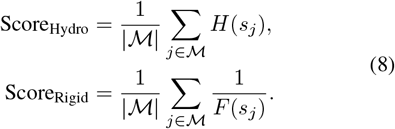

where *H*(*s*_*j*_) and *F* (*s*_*j*_) denote the hydrophobicity and flexibility value of amino acid *s*_*j*_.

### Affinity correlation analysis

After computing all scores, we evaluate their correlation with experimental binding affinities, expressed as −log(*K*_*D*_) or equivalent (e.g., −log(*IC*_50_)). Spearman’s rank correlation coefficient (*ρ*) and Kendall’s tau (*τ*) are computed to assess the predictive power and biological relevance of each scoring method. We also examine correlations between statistical similarity scores and log-likelihoods to probe whether generative models implicitly capture evolutionary constraints.

## 4. Empirical Evaluation

### 4.1. Datasets

In this study, we use fourteen datasets drawn from four sources: Absci HER2 (Shanehsazzadeh et al., 2023b), IgDesign (Shanehsazzadeh et al., 2023a), Nature (Porebski et al., 2024), and proprietary datasets from AstraZeneca (AZ).

#### Absci HER2

These datasets involve HCDR re-designs of Trastuzumab, an antibody that targets HER2. Sequence generation was performed using a two-step pipeline: first, machine learning models were used to predict HCDR loop structures conditioned on the HER2 backbone (PDB:1N8Z, Chain C), the Trastuzumab framework, and the known epitope; second, sequences were generated via inverse folding on the predicted structures. While HCDR3 lengths ranged from 9 to 17 residues, we focus on sequences with HCDR3 length 13, consistent with the native antibody. HCDR1 and HCDR2 were fixed at 8 residues. Binding affinities (*K*_*D*_) were measured using a FACS-based ACE assay. We analyze two datasets: (1) a “zero-shot binders” set, and (2) an SPR-validated “control” set containing both binders and non-binders.

#### IgDesign

This set includes seven antigen targets—FXI, IL36R, C5, TSLP, IL17A, ACVR2B, and TNFRSF9. Antibodies were designed by mutating either the HCDR3 alone or all three CDRs on the heavy chain. Each design library was synthesized and experimentally tested using SPR.

#### Nature

We also include datasets reported by Porebski et al. (2024), covering HER2, IL7, and HEL. Mutations in anti-HER2 are limited to HCDR3, while anti-IL7 involves LCDR1 and LCDR3. The HEL dataset consists of nanobodies with mutations across all three CDRs. Dataset sizes range from 19 to 38 sequences. We use *K*_*D*_ values for HER2 and HEL, and *IC*_50_ for IL7. For structure-based methods, parental structures were predicted using ImmuneBuilder2 (HER2), IgFold (IL7), and NanoBody-Builder2 (HEL) (Abanades et al., 2023; Ruffolo et al., 2023) as described in (Ucar et al., 2024).

**AZ** These proprietary datasets consist of two antibody libraries targeting separate antigens. The first target includes 24 variants, generated via rational design across four regions (HCDR1–3 and LCDR3). The second comprises 85 sequences drawn from three design strategies: two rationally designed libraries (one mutating heavy chain CDRs, the other light chain CDRs) and a third created using a machine learning model introducing changes across all six CDRs. Binding measurements are reported as *qAC*_50_ for Target-1 and *K*_*D*_ for Target-2. For models requiring structure, we use the corresponding crystal structures for both targets.

### 4.2. Results

#### Benchmarking predictive power across scoring methods

We assess the correlation between several scoring methods—including DiffAbXL-A log-likelihoods, BLOSUM similarity, PWM similarity, and PSSM similarity—and experimentally measured binding affinities across fourteen benchmark datasets. DiffAbXL-A was selected based on prior findings that its log-likelihood scores exhibit the strongest correlation with experimental binding affinity in (Ucar et al., 2024). Spearman’s rank correlation coefficients (*ρ*) are summarized in Table 1, and correlation statistics between log-likelihoods and BLOSUM or statistical similarity scores are shown in Table 2, with selected examples visualized in Figures 1 and 2 respectively. Log-likelihood scores derived from the DiffAbXL-A model in De Novo (DN) mode show consistent and often promising correlations with binding affinity across diverse design tasks. For example, we observe *ρ* = 0.43 on Absci HER2 Zero Shot, *ρ* = 0.62 on Nature HEL, and *ρ* = 0.62 on IgDesign IL17A. These results suggest that the model is capturing sequence features that are predictive of functional binding, even though it was not trained on the specific antibody libraries present in these datasets or on binding affinity prediction tasks. On the Nature IL7 dataset, where inhibition rather than binding affinity is measured (via *IC*_50_), a strong negative correlation is observed (*ρ* = −0.79). However, *IC*_50_ reflects the concentration needed to achieve 50% inhibition of a biological response, and is influenced by multiple factors beyond binding—such as receptor expression levels, signaling kinetics, and assay-specific artifacts. Unlike *K*_*D*_, which directly measures molecular interaction strength, *IC*_50_ integrates downstream effects and may vary substantially even when two molecules have similar affinities. As a result, correlations involving *IC*_50_ should be interpreted cautiously.

**Table 1.** Spearman correlations between binding affinity and scores from DiffAbXL-A-DN, biophysical features, PWM, PSSM, and BLOSUM. *, **, *** denote p-values ¡ 0.05, 0.01, and 1e-4, respectively. Affinity metrics are *q*AC50 for AZ Target-1, *IC*50 for IL7, and *K*_*D*_ for others.

| Approach | Model | Absci HER2 |  | Nature |  |  | AZ |  | Absci IgDesign |  |  |  |  |  |  |
| --- | --- | --- | --- | --- | --- | --- | --- | --- | --- | --- | --- | --- | --- | --- | --- |
|  |  | Zero Shot | Control | HEL | IL7 | HER2 | Target-1 | Target-2 | IL17A | ACVR2B | FXI | TNFRSF9 | IL36R | C5 | TSLP |
| Diffusion | DiffAbXL-A-DN | 0.43*** | 0.22*** | 0.62** | -0.79*** | 0.37* | -0.11 | 0.41** | 0.62** | 0.54* | 0.18 | 0.18 | 0.14 | -0.32 | -0.02 |
| Biophysical | Hydrophobicity | -0.17* | 0.04 | -0.13 | 0.57* | -0.38 | 0.30 | 0.14 | 0.49 | 0.49 | -0.39* | 0.41* | 0.70*** | 0.32 | -0.05 |
|  | Rigidity | 0.09 | 0.45*** | -0.18 | 0.33 | -0.37 | 0.18 | 0.06 | 0.49 | 0.49 | -0.22 | -0.16 | 0.15 | 0.19 | -0.12 |
| Statistical | PWM<sub>OAS</sub> | 0.30*** | 0.12* | 0.29 | 0.34 | 0.17 | -0.23 | 0.17 | -0.24 | 0.13 | -0.14 | 0.02 | -0.16 | -0.03 | -0.15 |
|  | PWM<sub>SAbDab</sub> | 0.30*** | 0.10* | 0.35* | -0.49* | 0.43* | 0.24 | 0.16 | -0.10 | 0.42 | -0.23 | 0.29 | -0.32* | -0.31 | 0.03 |
|  | PSSM<sub>OAS</sub> | 0.36*** | 0.22*** | 0.28 | -0.07 | 0.20 | -0.02 | 0.05 | -0.13 | 0.11 | -0.20 | 0.06 | -0.29 | -0.10 | -0.103 |
|  | PSSM<sub>SAbDab</sub> | 0.40*** | 0.29*** | 0.19 | -0.40 | 0.31 | 0.38 | 0.10 | -0.10 | 0.48* | -0.18 | 0.28 | -0.39* | -0.32 | -0.05 |
|  | PSSM<sub>DS</sub> | 0.50*** | 0.47*** | 0.52** | -0.18 | -0.16 | 0.29 | 0.24* | -0.01 | 0.24 | 0.53** | 0.05 | 0.26 | 0.00 | -0.11 |
|  | BLOSUM45<sub>G<sub>OAS</sub></sub> | 0.36*** | 0.03 | 0.26 | -0.59** | 0.26 | -0.52** | -0.07 | 0.10 | 0.03 | 0.29* | 0.03 | -0.27 | -0.25 | -0.36*** |
|  | BLOSUM45<sub>G<sub>SAbDab</sub></sub> | 0.39*** | 0.095 | 0.22 | -0.80*** | -0.34 | -0.21 | 0.16 | -0.39 | -0.006 | -0.42** | 0.26 | -0.27 | -0.32 | -0.34** |
|  | BLOSUM45<sub>DS</sub> | 0.46*** | 0.31*** | 0.52** | 0.69** | -0.32 | 0.43* | 0.30** | 0.01 | 0.47* | 0.52** | 0.17 | 0.38* | 0.10 | -0.29** |
|  | BLOSUM90<sub>PA</sub> | 0.48*** | 0.25*** | 0.57** | -0.86*** | -0.74*** | -0.09 | 0.26* | 0.77 | 0.63** | 0.32 | 0.41* | 0.57** | 0.57** | 0.19* |
|  | BLOSUM80<sub>PA</sub> | 0.48*** | 0.26*** | 0.58** | -0.85*** | -0.73*** | -0.10 | 0.24* | 0.77 | 0.73** | 0.32 | 0.42* | 0.58** | 0.64** | 0.20* |
|  | BLOSUM62<sub>PA</sub> | 0.48*** | 0.26*** | 0.57** | -0.85*** | -0.72*** | -0.06 | 0.26* | 0.77 | 0.71** | 0.32 | 0.38* | 0.58** | 0.60** | 0.21* |
|  | BLOSUM45<sub>PA</sub> | 0.50*** | 0.29*** | 0.59** | -0.87*** | -0.71*** | -0.08 | 0.30** | 0.77 | 0.73** | 0.33 | 0.48** | 0.50** | 0.62** | 0.22* |

**Table 2.**
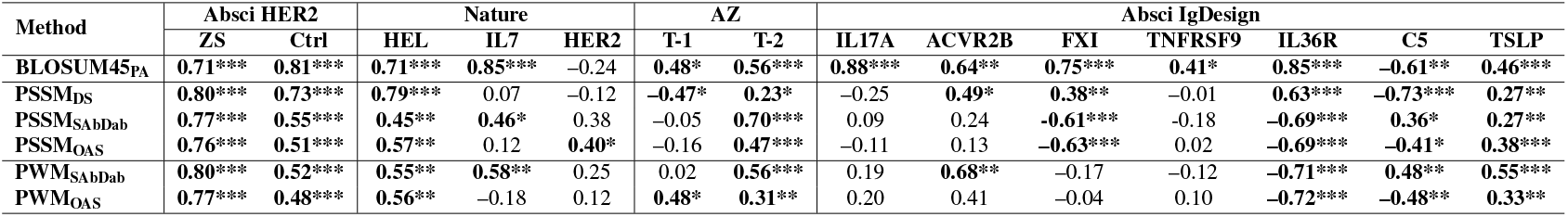
Spearman correlations between DiffAbXL-A-DN log-likelihoods and sequence similarity scores (BLOSUM45, PWM, PSSM). *, **, *** denote p-values ¡ 0.05, 0.01, and 1e-4, respectively.

**Figure 1.**
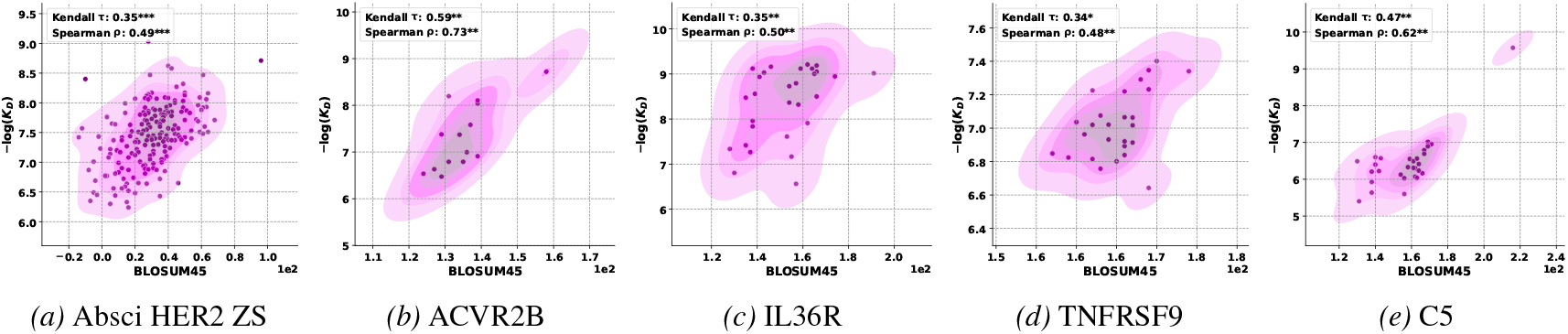
Correlation between BLOSUM45 and −*logK*_*D*_ : **a)** HER2 Zero-Shot (ZS), **b)** ACVR2B, **c)** IL36R, **d)** TNFRSF9, **e)** C5. *, **, *** indicate p-values under 0.05, 0.01 and 1e-4 respectively.

**Figure 2.**
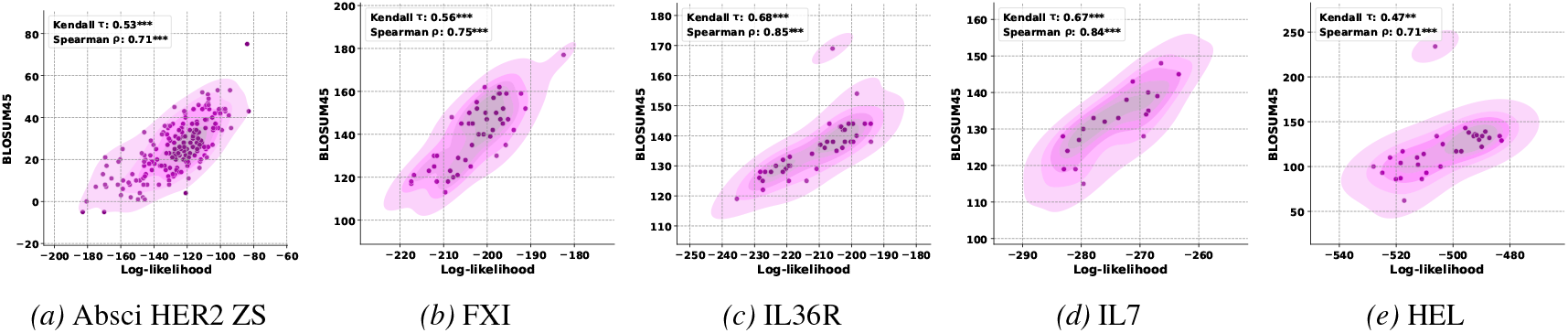
Correlation between BLOSUM45 and the log-likelihood scores of DiffAbXL-A-DN: **a)** HER2 Zero-Shot (ZS), **b)** FXI, **c)** IL36R, **d)** IL7, **e)** HEL. *, **, *** indicate p-values under 0.05, 0.01 and 1e-4 respectively.

### BLOSUM similarity scores align strongly with binding affinity

Across the board, BLOSUM-based similarity—particularly using BLOSUM45—shows good correlation with binding affinity. This trend holds across nearly all datasets, including Absci HER2, Nature HEL, and several IgDesign targets. For instance, BLOSUM45 achieves *ρ* = 0.50 on Absci HER2 Zero Shot, *ρ* = 0.59 on Nature HEL, and *ρ* = 0.73 on IgDesign ACVR2B. This finding supports the idea that evolutionary closeness to the parental (reference) sequence is a strong indicator of retained binding functionality. The consistent performance of BLOSUM matrices, especially those calibrated for more distant homologs (e.g., BLOSUM45), suggests they capture robust patterns relevant to antigen recognition and molecular stability. We note that across all datasets examined, the parental antibody—used as the reference for computing BLOSUM scores and thus assigned the highest similarity score by design—is consistently among the strongest binders. We expected that if this were not true, the correlation between BLOSUM similarity and binding affinity would be substantially weaker.

### Consensus-based BLOSUM scores lose predictive power

In contrast, when BLOSUM similarity is computed between designed sequences and global consensus sequences—either from OAS or SAbDab—the correlation with binding affinity largely disappears. For example, 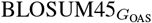 shows weak or negative correlations on many datasets, with significant drops observed on most datasets. This suggests that global evolutionary priors do not substitute well for dataset-specific reference sequences in affinity prediction.

### PWM similarity shows modest utility, with SAbDab out-performing OAS

PWM scores based on the OAS repertoire show weak and inconsistent correlation with binding affinity, with meaningful results limited to the Absci HER2 datasets (e.g., *ρ* = 0.30 on Zero Shot). However, when PWMs are computed from the SAbDab dataset, the correlation improves slightly. In addition to Absci HER2, we observe positive correlations on Nature HEL (*ρ* = 0.35) and Nature HER2 (*ρ* = 0.43), suggesting that structure-based databases such as SAbDab may better reflect the selective pressures acting on antibody binding regions since they contain antibody-antigen complexes. Nonetheless, performance remains below that of BLOSUM and DiffAbXL-A. This may be due to the local and repertoire-specific nature of the PWM used, which reflects background amino acid usage rather than target-specific substitution effects. Because PWMs are constructed from observed frequencies rather than substitution dynamics, they may fail to capture functionally relevant mutations, especially when applied outside their source distribution.

### PSSM similarity improves with data-specific priors

PSSM scores show variable performance depending on how the matrices are derived. Global PSSMs constructed from the OAS or SAbDab datasets yield modest correlations with binding affinity, with improvements observed for SAbDab-based PSSMs in some datasets (e.g., Absci HER2 and IgDesign ACVR2B). However, when PSSMs are constructed specifically from the dataset under evaluation (**PSSM**_**DS**_), correlation improves substantially. For example, **PSSM**_**DS**_ yields *ρ* = 0.50 on Absci HER2 Zero Shot and *ρ* = 0.52 on Nature HEL, both surpassing the performance of global PSSMs. These results highlight the importance of local sequence context in capturing meaningful constraints for affinity prediction and suggest that dataset-specific PSSMs can serve as useful tools when sufficient in-distribution sequence data are available.

### Log-likelihood scores correlate with BLOSUM, PSSM, and PWM similarity metrics

We observe strong correlations between DiffAbXL-A log-likelihood scores and BLOSUM45 similarity across most datasets (Table 2), with 12 out of 14 cases exceeding a Spearman *ρ* of 0.4, and several surpassing 0.8 (e.g., IL17A, IL36R, IL7). This suggests that the generative model is implicitly learning amino acid substitution patterns that align closely with established evolutionary priors. Similar, though generally weaker, correlations are observed with PWM-based scores. Notably, PWMs derived from SAbDab show stronger alignment with log-likelihoods than those from OAS, especially in datasets such as IL7 (*ρ* = 0.58) and ACVR2B (*ρ* = 0.68). PSSM similarity also correlates with model scores, particularly when computed from dataset-specific (**PSSM**_**DS**_) or SAbDab-based matrices, supporting the view that the model internalizes substitution preferences.

### Biophysical scores show target-specific utility

Two coarse-grained biophysical metrics—Kyte-Doolittle hydrophobicity and Karplus-Schulz rigidity—were also evaluated. Hydrophobicity content correlates positively with binding affinity on Nature IL7 (*ρ* = 0.57) and IgDesign IL36R (*ρ* = 0.70), suggesting a potential link between hydrophobic residues and binding affinity in these datasets. Rigidity shows a moderate correlation only on Absci HER2 Control (*ρ* = 0.45), with little signal elsewhere. These results indicate that such physicochemical scores may capture some target-specific trends but are not general-purpose predictors of affinity.

### Consistent evolutionary signatures in log-likelihoods from diverse generative models

As with DiffAbXL-A, log-likelihoods from both AntiFold and IgLM show strong correlations with BLOSUM45 similarity scores (Table 3). For example, AntiFoldPA log-likelihoods correlate with BLOSUM similarity at *ρ* = 0.76 on Nature HEL and *ρ* = 0.80 on IgDesign TSLP. Similarly, IgLM[bi]_PA_ achieves *ρ* = 0.73 on Nature HEL and shows moderate-to-strong alignment on multiple other datasets. These findings further support the hypothesis that generative models implicitly learn substitution preferences that align with classical evolutionary priors. We also note that when log-likelihoods (LLs) correlate with binding affinity (*K*_*D*_), they almost always exhibit an even stronger correlation with BLOSUM similarity. However, the converse does not necessarily hold: a strong LL–BLOSUM correlation does not imply a meaningful LL–*K*_*D*_ correlation.

**Table 3.** Spearman correlations between model log-likelihoods (LLs) and *K*_*D*_ values, as well as LLs and BLOSUM45_PA_ scores, for DiffAbXL-A-DN, AntiFold, and IgLM. *, **, *** denote p-values ¡ 0.05, 0.01, and 1e-4, respectively.

| Model | Metric | Absci HER2 |  | Nature |  | Absci IgDesign |  |  |  |  |  |  |
| --- | --- | --- | --- | --- | --- | --- | --- | --- | --- | --- | --- | --- |
|  |  | ZS | Ctrl | HEL | HER2 | IL17A | ACVR2B | FXI | TNFRSF9 | IL36R | C5 | TSLP |
| DiffAbXL-A-DN | LL vs $K_D$ | 0.43*** | 0.22*** | 0.62** | 0.37* | 0.62** | 0.54* | 0.18 | 0.18 | 0.14 | -0.32 | -0.02 |
|  | LL vs BLOSUM45<sub>PA</sub> | 0.71*** | 0.81*** | 0.71*** | -0.24 | 0.88*** | 0.64** | 0.75*** | 0.41* | 0.85*** | -0.61** | 0.46*** |
| AntiFold<sub>PA</sub> | LL vs $K_D$ | 0.45*** | 0.15** | 0.64*** | -0.38 | -0.79** | 0.29 | 0.02 | 0.04 | 0.18 | -0.01 | 0.24** |
|  | LL vs BLOSUM45<sub>PA</sub> | 0.61*** | 0.61*** | 0.76*** | 0.60** | -0.39 | -0.04 | 0.14 | 0.65*** | -0.22 | 0.01 | 0.80*** |
| IgLM[pre]<sub>PA</sub> | LL vs $K_D$ | -0.09 | 0.22*** | 0.35* | 0.02 | 0.12 | 0.16 | 0.31* | 0.12 | -0.63*** | -0.26 | 0.23* |
|  | LL vs BLOSUM45<sub>PA</sub> | -0.22** | -0.07 | 0.57** | -0.33 | 0.16 | -0.36 | 0.60*** | 0.11 | -0.43** | -0.82*** | 0.51*** |
| IgLM[bi]<sub>PA</sub> | LL vs $K_D$ | 0.26** | -0.05 | 0.59** | -0.12 | 0.14 | -0.01 | -0.11 | -0.09 | 0.24 | -0.21 | -0.08 |
|  | LL vs BLOSUM45<sub>PA</sub> | 0.35*** | 0.14** | 0.73*** | 0.43* | 0.56* | 0.48* | -0.58*** | 0.38* | -0.04 | -0.59** | 0.36*** |

**Table 4.**
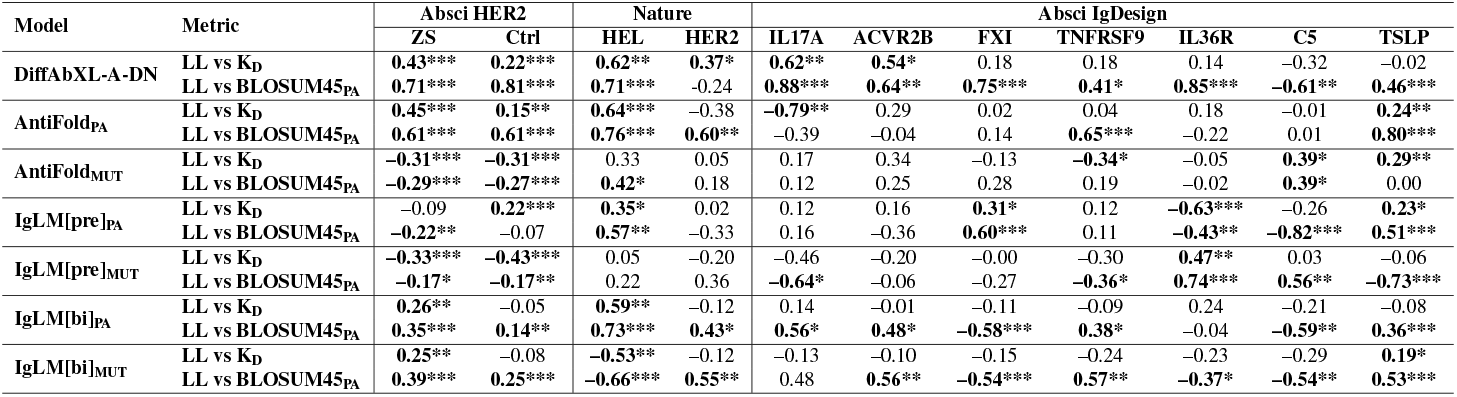
Spearman correlations between the model log-likelihoods (LLs) and K_D_ values as well as between LLs and BLOSUM45_PA_ scores for DiffAbXL-A-DN, AntiFold and IgLM. *, **, *** indicate p-values under 0.05, 0.01, and 1e-4, respectively.

#### 4.2.1. Large-scale HER2 screening using deep-screening intensities

To complement our affinity-based benchmarks, we analyze the full HER2 deep-screening dataset from Porebski et al. (2024), which reports high-throughput fluorescence intensities (FIs) as proxies for antibody binding. After removing sequencing artifacts, we retain approximately 179k unique HCDR3 variants, each associated with a mean FI aggregated across replicate measurements. Following the threshold used in the authors’ repository (Holliger Lab, 2024), we define strong binders as sequences with FI ≥ 250. The parental (WT) sequence, by contrast, exhibits an FI of 136.31 (Fig. 3a), below the strong binder threshold.

**Figure 3.**
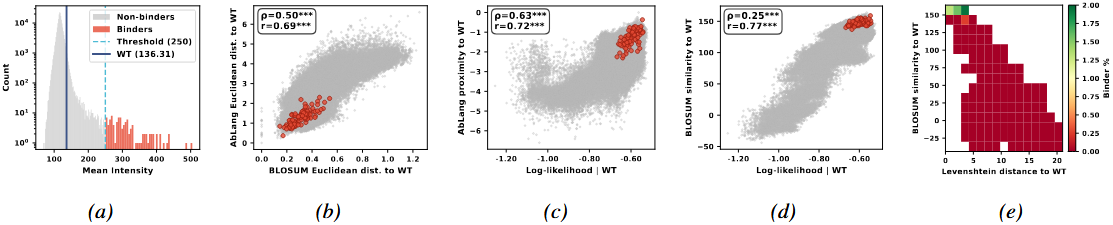
Large-scale HER2 screening. **a)** Distribution of antibody intensities from deep screening; WT has FI = 136.31. **b)** Correlation between distances to WT in AbLang-2 and BLOSUM45 embedding spaces. c) Correlation between AbLang-2 proximity to WT (inverse embedding distance) and log-likelihood. d) Correlation between BLOSUM45 similarity to WT and log-likelihood. e) Hit-rate heatmap showing strong-binder enrichment by BLOSUM similarity and edit distance; enrichment peaks at BLOSUM ≥ 150 and Levenshtein ≤ 4. Significance: * p < 0.05, ** p < 0.01, *** p < 1e-4.

To assess how language models and classical similarity metrics relate to binding in this large library, we compute each variant’s distance to WT in AbLang-2 embedding space (Olsen et al., 2024) and in a BLOSUM45-based substitution space (Gessner et al., 2024). These distances are strongly correlated (Spearman *ρ* = 0.50, Pearson *r* = 0.69; Fig. 3b), indicating that AbLang-2’s learned representation aligns closely with BLOSUM-derived substitution patterns.

We next examine how these distances relate to likelihood scores. AbLang-2 log-likelihoods correlate strongly with proximity to WT in its embedding space (Spearman *ρ* = 0.63, Pearson *r* = 0.72; Fig. 3c), and also with BLOSUM45 similarity (Spearman *ρ* = 0.25, Pearson *r* = 0.77; Fig. 3d). While the rank correlation is weaker for BLOSUM, both relationships suggest that high-likelihood sequences tend to remain evolutionarily close to WT.

Finally, we assess where strong binders concentrate in sequence space. A density heatmap over BLOSUM45 similarity and Levenshtein distance to WT (Fig. 3e) reveals that strong binders are most enriched in regions with BLOSUM similarity ≥ 150 and edit distance ≤ 4. While both metrics capture local enrichment, BLOSUM similarity provides a clearer separation between binders and non-binders in this setting, suggesting it is a more informative proxy for WT proximity in large-scale screening.

## 5. Conclusion

We examined the connection between generative model scores and classical evolutionary similarity metrics in the context of antibody design. Across fourteen datasets of experimentally characterized antibody variants, BLOSUM similarity to the wild type—particularly using matrices such as BLOSUM45—showed strong and consistent correlation with binding affinity, often rivaling or exceeding the performance of model-based log-likelihood scores. Among generative models, the diffusion-based model DiffAbXL-A showed the highest correlation with affinity values, consistent with prior findings in (Ucar et al., 2024). Moreover, evaluations of AntiFold and IgLM revealed that their log-likelihood scores also align with BLOSUM similarities, even when their correlation with binding affinity is weaker. This reinforces the idea that generative models may implicitly learn substitution patterns shaped by evolutionary pressure, even without explicit supervision.

In contrast, PWM-based scores exhibited limited and variable performance, particularly when derived from general antibody repertoires. Slight improvements were observed with PWMs constructed from antibodies in complex with antigens, though these still underperformed relative to BLOSUM similarities over the parental antibody sequence in each dataset. PSSM-based scores showed somewhat stronger and more stable correlations than PWMs, especially when constructed from dataset-specific alignments, occasionally matching the predictive power of BLOSUM and log-likelihood scores. However, their performance was less consistent when based on broad repertoires such as OAS or structure databases such as SAbDab. Simple biophysical descriptors such as hydrophobicity and rigidity captured some signal in a few cases but lacked generalizability.

These findings highlight the utility of interpretable, evolution-derived metrics such as BLOSUM and underscore the importance of contextual information—such as reference sequence choice—in score interpretation. They also suggest that generative models encode evolutionary signals that can be leveraged for scoring and prioritization. Large-scale HER2 screening further validates that WT-referenced BLOSUM similarity and language-model likelihoods behave as concordant distance-to-binder signals, providing a mechanistic explanation for their correlation with binding. As generative models continue to improve, understanding the extent to which their representations align with biological priors will be essential for advancing robust and interpretable generative models for therapeutic antibody design.

## Impact Statement

This research advances the use of machine learning in antibody engineering by investigating the predictive power of statistical models such as BLOSUM and PWM for ranking antibody designs based on binding affinity as well as studying their relationship with the log-likelihood scores of diffusion-based generative models. By demonstrating an association between derived scores and real-world experimental data, this work provides a pathway to accelerate therapeutic antibody discovery while minimizing costly trial-and-error experimentation. Ethical and societal impacts, such as improved healthcare outcomes and broader accessibility of life-saving treatments, mirror the established considerations in the broader field of machine learning-driven drug discovery.

## Appendix

### A. License Information

IgDesign datasets (Shanehsazzadeh et al., 2023a) are released under MIT license. Absci Her2 datasets (Shanehsazzadeh et al., 2023b) are released under BSD License. SAbDab and OAS datasets are available under a CC-BY 4.0 license. We will release our code upon the acceptance of our paper with Apache 2.0 license.

#### B. Construction of PWM and PSSM

To analyze amino acid preferences at each structurally equivalent position, we first aligned all sequences using the AHo numbering scheme, which provides a consistent positional framework across antibody variable domains. Each sequence was mapped into an AHo-labeled position–residue dictionary, and alignment matrices were constructed accordingly.

We computed the Position Weight Matrix (PWM) by counting the occurrences of each amino acid (including gaps) at each AHo-defined position across all aligned sequences. To prevent zero probabilities and ensure numerical stability, a Laplace pseudocount of 1 was added to each residue count. The resulting frequency *f*_*i,a*_ of amino acid *a* at position *i* was computed as:

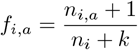

where *n*_*i,a*_ is the count of amino acid *a* at position *i, n*_*i*_ is the total number of observations at that position (including gaps), and *k* = 21 is the number of possible residue types (20 amino acids plus the gap character). The PWM thus represents a normalized probability distribution over residues at each position.

To construct the Position-Specific Scoring Matrix (PSSM), we first estimated background frequencies *q*_*a*_ for each amino acid *a* across the entire aligned dataset, again applying Laplace pseudocounts:

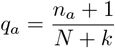

where *n*_*a*_ is the total count of amino acid *a* in the full alignment, *N* is the total number of observed residues across all positions and sequences (including gaps), and *k* = 21 as before.

The log-odds score *S*_*i,a*_ for each residue *a* at position *i* was then calculated as:

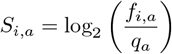

This score reflects how much more (or less) likely a residue is to appear at a specific position compared to its global background expectation. The final PSSM was stored as a position-by-residue matrix of log-odds scores. To summarize the most likely residue at each position, a consensus sequence was derived by selecting the amino acid with the highest frequency in the PWM.

### C. Additional Results

### D. Log-Likelihood Scoring

#### D.1. Setup and Notation

Let an antibody sequence be denoted by

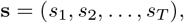

where each *s*_*t*_ represents one of the 20 canonical amino acids. A left-to-right autoregressive language model (e.g., GPT-2) defines the sequence probability via the factorization

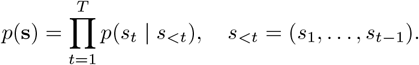

Let ℓ_*t*_( ·) ∈ ℝ^20^ denote the unnormalized logits output by the model at position *t*. The corresponding conditional log-probability is computed as

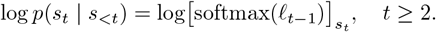

For *t* = 1, either a special start-of-sequence token or a uniform prior may be used.

#### D.2. Full Sequence Log-Likelihood

The total log-likelihood of a sequence **s** is given by

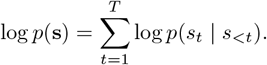

This quantity can be computed from a single forward pass through the model. The following pseudocode illustrates the computation using `transformers`-style APIs:

~~~
# Tokenize and run one forward pass
input_ids = tokenizer.encode(seq)               # shape [1, T]
logits = model(input_ids).logits                # shape [1, T, 20]
log_probs = softmax(logits, dim=-1).log()
# Shift so that log_probs[*, t-1, *] = log p(s_t | s_<t)
shifted = log_probs[:, :-1, :]
labels = input_ids[:, 1:]
# Gather and sum
token_ll = shifted.gather(-1, labels.unsqueeze(-1)).squeeze(-1)
total_ll = token_ll.sum()
~~~

#### D.3. Single Contiguous Masked Region

To compute the log-likelihood of a contiguous span (*s*_*a*_, …, *s*_*b*_) conditioned on its left context, we can use

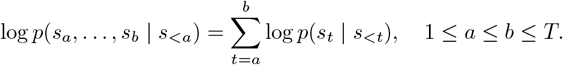

Given the vector of log-probabilities computed above, the relevant entries are summed as follows:

~~~
# token_ll[i] = log p(s_{i+1} | s_<=i)
span_ll = token_ll[(a - 1) : b].sum()
~~~

#### D.4. Multiple Disjoint Masked Regions

Consider a collection of *K* disjoint spans 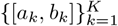. The total log-likelihood over these regions is

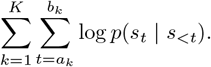

This is equivalent to summing over the union of all token positions in the selected spans:

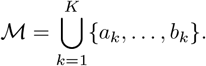

~~~
positions = []
for (a, b) in spans:
     positions.extend(range(a - 1, b))
multi_ll = token_ll[positions].sum()
~~~

#### D.5. Context Choice for Library Scoring: LL_MUT_ vs. LL_PA_

For a designed antibody library derived from a common parental sequence, the log-likelihood of each designed sequence can be computed using one of two distinct approaches:

##### Mutation-context likelihood (LL_MUT_)

In this setting, the model is conditioned directly on each designed (mutant) sequence to compute its own log-likelihood:

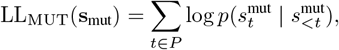

where *P* denotes the set of mutated positions. This approach reflects the model’s confidence in the mutant sequence given the full autoregressive context of the design.

##### Parent-context likelihood (LL_PA_)

Alternatively, model logits may be obtained from the original parental sequence **s**_pa_, and the log-likelihood is then evaluated using the designed sequence **s**_mut_ by gathering log-probabilities only at mutated positions:

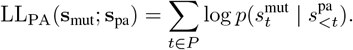

In practice, the parental sequence is passed to the model to obtain the sequence of conditional distributions, and the designed sequence is used only to select which token probabilities to score at positions *t* ∈ *P*.

##### Implementation notes

Let `input pa`denote the tokenized parental sequence, and `input mut`denote the mutant sequence. Then:

~~~
# For LL_MUT
logits_mut = model(input_mut).logits log_probs_mut = softmax(logits_mut, dim=-1).log()
# Use positions P on token_ll_mut to compute LL_MUT
# For LL_PA
logits_pa = model(input_pa).logits
log_probs_pa = softmax(logits_pa, dim=-1).log() #
Gather log_probs_pa at positions t in P,
# using tokens from input_mut for indexing
~~~

##### Use cases

LL_MUT_ reflects how likely a full designed sequence is under the model’s distribution, incorporating all mutated residues and their autoregressive influence. In contrast, LL_PA_ measures how well the mutated residues are supported by the local sequence context inherited from the parent, isolating the evaluation to mutation sites without considering their downstream impact.

### Summary

- LL_MUT_: evaluates mutant sequence under its own autoregressive context.
- LL_PA_: evaluates mutant residues in the fixed parental context.

### E. IgLM Log-Likelihood Scoring

#### E.1 Preceding vs. Bidirectional Context Scoring

IgLM is a decoder-only Transformer model trained with an infilling objective designed for antibody sequence modeling (Shuai et al., 2023). Instead of standard left-to-right language modeling, IgLM is trained to reconstruct masked spans within a sequence using both left and right flanking context. During training, a contiguous span of amino acids is removed and replaced with a special `[MASK]`token. The remaining prefix and suffix of the sequence are concatenated, separated by a `[SEP]`token, and the removed span is appended after this context, followed by an `[ANS]`token to indicate the end of the span. This reordered sequence is used as input to the model, which is then trained to autoregressively predict the span tokens (and the `[ANS]`terminator), conditioned on the entire flanking context.

Formally, for a sequence **s** = (*s*_1_, …, *s*_*T*_) with a masked span *S* = (*s*_*s*_, …, *s*_*e*_), IgLM constructs an input of the form:

[CHAIN] [SPECIES]*s*_1_, …, *s*_*s*−1_, `[MASK]`, *s*_*e*+1_, …, *s*_*T*_, `[SEP]`, *s*_*s*_, …, *s*_*e*_, `[ANS]` where `[CHAIN]`and `[SPECIES]`are fixed metadata tokens indicating chain type and species. During evaluation, IgLM supports two log-likelihood computation strategies:

- Preceding context (autoregressive) scoring ([pre]): left-to-right log-likelihoods are computed over the full sequence.
- Bidirectional context (infilling-based) scoring ([bi]): log-likelihoods are computed using bidirectional context, consistent with the training setup.

Both strategies can be used with either the mutant or parental sequence as input context. For example, `IgLM[pre]`_MUT_ uses the mutant sequence for autoregressive scoring, while `IgLM[bi]`_MUT_ uses the mutant sequence for bidirectional infilling.

##### Preceding-Context Scoring

In the [pre] setting, the full sequence (mutant or parental) is passed to the model, and token-level log-likelihoods are computed left-to-right. For a given context sequence **s**^CTX^, and mutant target **s**^mut^, the log-likelihood is:

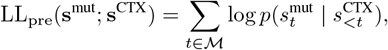

where ℳ is the set of mutated positions, and CTX ∈ {MUT, PA} indicates the context source.

##### Bidirectional (Infilling-Based) Scoring

In the [bi] setting, IgLM uses its span-infilling mechanism to evaluate masked spans with bidirectional context. For each mutated span 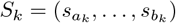, the span is masked in the context sequence, and the corresponding mutant residues are passed as the target:

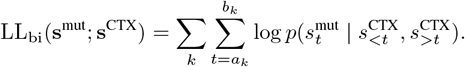

As before, CTX indicates whether the context is the parental or mutant sequence.

##### Implementation Considerations

The IgLM implementation supports both evaluation protocols. For autoregressive scoring, the model processes the entire context sequence once and gathers log-probabilities at mutation sites. For infilling-based scoring, the model must be called separately for each mutated span: the corresponding region is masked in the context sequence, and the mutant tokens are appended as the infill segment. The model then computes log-probabilities for these tokens after the `[SEP]`marker, consistent with its training objective.

As IgLM was trained to model masked spans using bidirectional context, the infilling-based scoring ([bi]) is more aligned with the model’s architecture and is empirically reported to yield lower perplexity than autoregressive scoring. However, both modes are supported and yield useful comparisons when paired with either mutant or parental input.

##### Summary

- LL_pre_: computes log-likelihood using the full sequence as input, relying only on autoregressive left context.
- LL_bi_: computes log-likelihood of the designed regions using IgLM’s infilling mechanism, conditioned on bidirectional context from the sequence.

### F. AntiFold Log-Likelihood Scoring

AntiFold is an antibody-specific inverse folding model based on the ESM-IF1 architecture, trained to predict sequences given fixed backbone structures (Høie et al., 2023). Given a structure input, AntiFold generates amino acid sequences autoregressively from N-to C-terminus using a decoder-only Transformer architecture with *causal attention*. This means each position attends only to its preceding sequence positions and not to future residues. However, AntiFold conditions globally on the full backbone structure, which is processed separately and fed as a contextual embedding at each decoding step.

During evaluation, AntiFold outputs the log-probability assigned to each of the 20 amino acids at each sequence position. These per-position log-probabilities can be used to compute the log-likelihood of any specified subset of residues, including disjoint masked regions.

#### Mutation-context likelihood (LL_MUT_)

In the first evaluation strategy, the model is run using the full mutant sequence as input, along with the associated structure^1^. Per-position log-probabilities are extracted from the model output, and the log-likelihood of the mutant residues is computed over the specified masked positions:

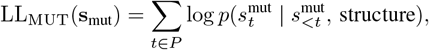

where *P* denotes the indices of mutated residues. Because the input sequence is the mutant itself, the decoder conditions on the correct mutated left context for each position in *P*.

##### Parent-context likelihood (LL_PA_)

Alternatively, log-probabilities can be computed using the parental sequence as input. In this case, the model is conditioned on the original (pre-mutation) left context, and the mutant amino acids are scored using the model’s output:

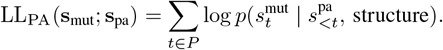

This formulation assesses how well the mutant residues fit into the structural and sequential context defined by the parent. However, since the decoder is causal, any mutations occurring at early positions can affect the correctness of conditioning for later positions if the full mutant context is not used.

##### Implementation details

AntiFold outputs a matrix of per-position log-probabilities in CSV format. Given a mutant sequence and set of mutated positions *P*, the log-likelihood is computed by gathering the model’s log-probability for each mutant residue at the corresponding position:

~~~
# For LL_MUT
log_probs_mut = antifold(model_input=mutant.pdb)
ll_mut = sum([log_probs_mut[t][s_mut[t]] for t in P])
# For LL_PA
log_probs_pa = antifold(model_input=parent.pdb)
ll_pa = sum([log_probs_pa[t][s_mut[t]] for t in P])
~~~

Here, `s mut[t]`refers to the amino acid at position *t* in the mutant sequence, and `log probs[t][aa]`gives the log-probability assigned to amino acid `aa`at position *t*.

##### Summary

- LL_MUT_: scores the mutant using its own autoregressive context.
- LL_PA_: scores the mutant residues using log-probabilities from the parent context.
- In both cases, log-likelihood is computed by summing over specified mutated regions.

## Footnotes

1 Backbone structure of mutant sequence is assumed to be same as the parental sequence

